# *PTBP1* mRNA isoforms and regulation of their translation

**DOI:** 10.1101/509174

**Authors:** Luisa M. Arake de Tacca, Mia C. Pulos, Stephen N. Floor, Jamie H. D. Cate

## Abstract

Polypyrimidine tract-binding proteins (PTBPs) are RNA binding proteins that regulate a number of post-transcriptional events. Human PTBP1 transits between the nucleus and cytoplasm and is thought to regulate RNA processes in both. However, information about *PTBP1* mRNA isoforms and regulation of PTPB1 expression remain incomplete. Here we mapped the major *PTBP1* mRNA isoforms in HEK293T cells, and identified alternative 5’ and 3’untranslated regions (5’ UTRs, 3’ UTRs) as well as alternative splicing patterns in the protein coding region. We also assessed how the observed *PTBP1* mRNA isoforms contribute to PTBP1 expression in different phases of the cell cycle. Previously, *PTBP1* mRNAs were shown to crosslink to eukaryotic translation initiation factor 3 (eIF3). We find that eIF3 binds differently to each *PTBP1* mRNA isoform in a cell cycle-dependent manner. We also observe a strong correlation between eIF3 binding to *PTBP1* mRNAs and repression of PTBP1 levels during the S phase of the cell cycle. Our results provide evidence of translational regulation of PTBP1 protein levels during the cell cycle, which may affect downstream regulation of alternative splicing and translation mediated by PTBP1 protein isoforms.

## INTRODUCTION

PTBP1 was first discovered as a purified protein that bound to polypyrimidine tract regions of introns (Garcia-Blanco et al. 1989). Initially, PTBP1 was thought to be part of the splicing machinery, until U2AF65 was discovered as the splicing factor responsible for recognizing the polyU tracts at the 3’ splice site during the assembly of the spliceosome (Gil et al. 1991). PTBP1 has since been shown to regulate alternative exon selection during mRNA processing by repressing exon inclusion (Xue et al. 2009). Although PTBP1 acts as an alternative splicing (AS) factor in the nucleus, it also shuttles between the nucleus and cytoplasm. When PTBP1 is present in the cytoplasm, it is thought to be involved in post-transcriptional regulation, processes that require cap-independent translational control, RNA localization or changes in mRNA stability (Kamath et al. 2001; Romanelli et al. 2013). In addition to its role in molecular processes including splicing, polyadenylation, translation initiation, and mRNA stability, PTBP1 has recently been linked to the regulation of the cell cycle (Monzón-Casanova et al. 2018).

PTBP1 is a 57 kDa protein comprised of four RNA recognition motifs (RRMs) with a bipartite nuclear localization domain (NLD) and a nuclear export signal (NES) at the N-terminus of the protein (Li and Yen 2002; Pérez et al. 1997; Wollerton et al. 2001). The expression of PTBP1 is tightly regulated through alternative splicing events (Wollerton et al. 2004). Its 15 exons have previously been shown to be alternatively spliced into three major mRNA isoforms, termed *PTBP1-1, PTBP1-2*, and *PTBP1-4*. The first described isoform, *PTBP1-1* encodes a protein of 521 amino acids containing all four RRMs. The alternatively spliced isoforms, *PTBP1-2* and *PTBP1-4*, encode an additional 19 or 26 amino acids, respectively, between the RRM2 and RRM3 domains derived from exon 9 inclusion (Garcia-Blanco et al. 1989; Valcárcel and Gebauer 1997;Sawicka et al. 2008; Romanelli et al. 2013). Despite being very similar, the different isoforms have distinct roles in splicing and internal ribosome entry site (IRES)-mediated initiation of translation. The absence or length of the unstructured region between RRM2 and RRM3 results in differential recognition of target RNAs. These functional differences coupled with differing PTBP1 isoform ratios in different cell lines suggests that changes in relative PTBP1 isoform expression levels may be a cellular determinant of alternative splicing events (Gueroussov et al. 2015,Wollerton et al. 2001). For example, in the case of tropomyosin alternative splicing, PTBP1-4 represses exon 3 inclusion more than PTBP1-1 both *in vivo* and *in vitro*, whereas PTBP1-2 harbors intermediate activity (Wollerton et al. 2001). Additionally, differences in exon 9 skipping in *PTBP1* mRNAs has been found to affect the levels of many additional alternative splicing (AS) events, likely modulating the timing of transitions in the production of neural progenitors and mature neurons so as to affect brain morphology and complexity (Gueroussov et al. 2015).

In eukaryotic mRNAs, the 5’ and 3’untranslated regions (5’ and 3’ UTRs) serve as major *cis*-regulatory control elements. RNA sequences and structures in the 5’ UTR and 3’ UTR can act as binding sites for translation initiation factors and other RNA binding proteins to influence the translational output of an mRNA and its lifetime in the cell. To date, how the alternatively spliced isoforms of PTBP1 are connected to different 5’ UTRs and 3’ UTRs in *PTBP1* mRNA has not been determined. Several annotation databases, such as ENSEMBL (Ensembl Release 94) (Zerbino et al. 2018), FANTOM5 (Riken Center for Integrative Medical Sciences (IMS)) (Noguchi et al. 2017) and NCBI Gene (O’Leary et al. 2015), have information on *PTBP1* isoforms. However, the information on UTRs differs across these databases. In ENSEMBL, the three main isoforms have distinct 5’ UTRs and a common 3’ UTR. In the NCBI Gene (refseq) database, *PTBP1* has common 5’ and 3’ UTRs. The FANTOM5 database (The FANTOM Consortium and the RIKEN PMI and CLST, 2014) only accounts for two distinct 5’ UTRs for *PTBP1* and a common 3’ UTR. Finally, the APASdb database for polyadenylation signals (You et al. 2015) reports two major polyadenylation sites within the *PTBP1* 3’ UTR. These libraries need to be reconciled into a comprehensive model of *PTBP1* transcript isoforms allowing further biochemical analysis of the regulatory pathways that influence *PTBP1* mRNA isoform production and translation.

To better understand the regulation of *PTBP1* mRNA isoform levels in the cell, we mapped the major *PTBP1* mRNA variants present in mammalian HEK293T cells. We analyzed the 5’ UTR elements using 5’-RACE (RLM-RACE) and long-read sequencing (Oxford nanopore). We also mapped the 3’ UTRs and open reading frames. Using Western blots and mRNA reporters, we determined how the *PTBP1* mRNA isoforms are translated in different stages of the cell cycle. Previous evidence revealed that human translation initiation factor eIF3, the largest translation initiation factor, crosslinks to the 5’ UTR elements of several messenger RNAs, including *PTBP1*. While bound to mRNAs, eIF3 acts to either activate or repress their translation (Lee et al. 2015). For this reason, we also probed eIF3 interactions with *PTBP1* mRNAs to determine whether eIF3 may act as a *trans-acting* factor regulating *PTBP1* isoform translation.

## RESULTS

### Endogenous levels of PTBP1

Since PTBP1 has been implicated in regulating numerous processes including the cell cycle, we analyzed the endogenous levels of PTBP1 in HEK293T cells harvested in different stages of the cell cycle (Figure 1A). We observed that PTBP1 isoforms vary dramatically during cell cycle progression. Cells arrested in the G2 or M phases had the highest levels of all three isoforms (PTBP1-1, PTBP1-2, PTBP-4, Figure 1B), with the upper band, comprising PTBP1-2 and PTBP1-4 (Wollerton et al.2001), having a higher expression profile than PTBP1-1 regardless of cell cycle phase. All three isoforms exist at low levels in G1, and increase slightly in S, before a larger burst in G2/M occurs. Notably, *PTBP1* mRNA levels do not fluctuate as much as protein levels in the different stages of the cell cycle (Figure 1C). Although we did not separate the contributions of translation and protein degradation, these results indicate that post-transcriptional regulation of PTBP1 expression occurs as a function of the cell cycle.

**Figure 1.**
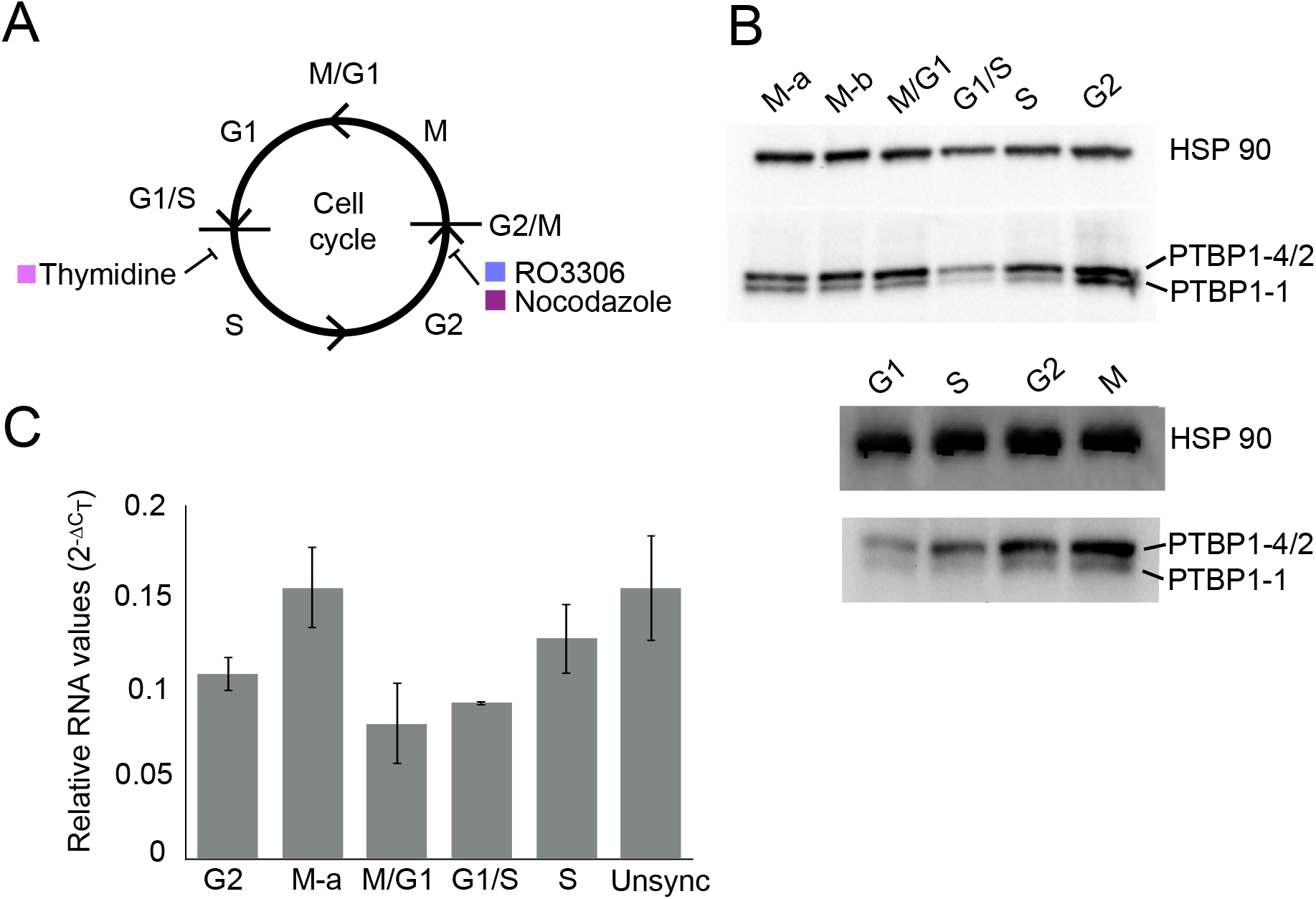
*PTBP1* expression changes across the cell cycle. **A.** Chemical inhibitors used to arrest cells at specific phases of the cell cycle: thymidine, arrest at G1/S, and RO3306 or Nocodazole, arrest at G2/M transitions. Cells were synchronized and collected at time points after release from the drugs. **B.** Western blots of whole cell lysates of synchronized HEK293T cells prepared using synchronized samples. Separate methods to arrest G2/M: M-a was synchronized with the use of RO3306, and M-b was synchronized with the use of Nocodazole. PTBP1-2 and PTBP1-4 protein isoforms have similar sizes and co-migrate in the gel. Bottom, independent biological replicate of the Western using RO3306. **C.** Total RNA amounts of each transcript were assessed using quantitative PCR for each phase of the cell cycle.

### Mapping the 5’ UTR, CDS and 3’ UTR sequences in *PTBP1* mRNAs

To test whether *PTBP1* transcript isoform sequences in the ENSEMBL database are in agreement with the transcription start sites (TSS) in FANTOM5, we employed RNA Ligase Mediated Rapid Amplification of cDNA Ends (RLM-RACE) and Nanopore sequencing of mRNAs extracted from HEK293T cells to map *PTBP1* transcripts (Figure 2). Although both TSS in the FANTOM5 database were confirmed by RLM-RACE, we could not verify the presence of the 5’ UTR for ENSEMBL transcript <u>ENST00000356948.10</u>. Notably, our RLM-RACE data supports a different TSS for ENSEMBL transcript <u>ENST00000349038.8</u>, 7 nucleotides (nts) 5’ of the annotated TSS, in agreement with the TSS mapped in the FANTOM5 database (Figure 2). The longer TSS for this transcript is also in agreement with the fact that eIF3 crosslinks to nucleotides 5’ of the ENSEMBL-annotated TSS (Figure 2, Figure 3).

**Figure 2.**
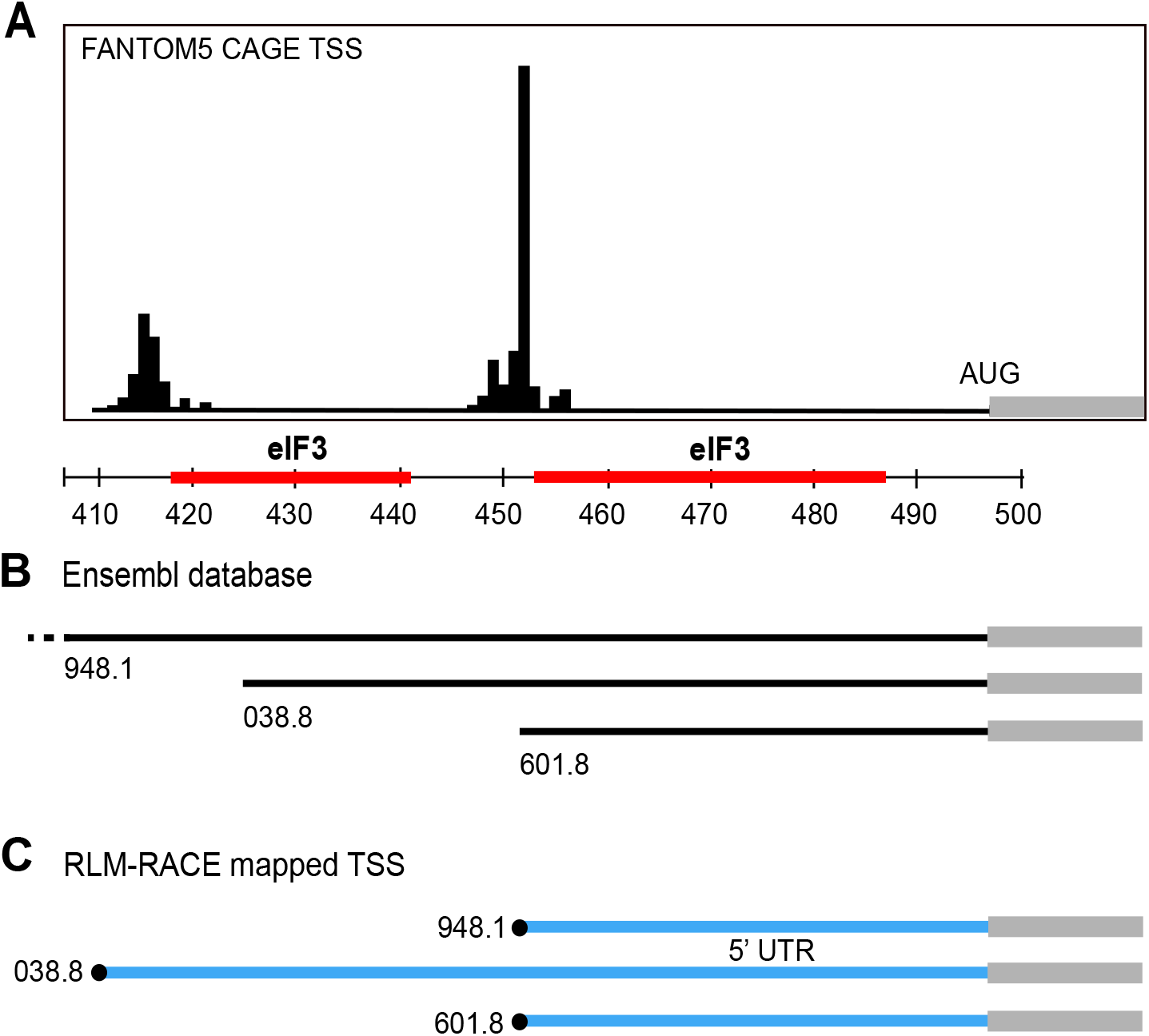
Database and experimental mapping of the three major transcript isoforms of *PTBP1* mRNA. **A.** Transcription start sites (TSS) determined by CAGE mapping (FANTOM5), aligned to the consensus PAR-CLIP crosslinking sites to eIF3 (red), along with hg38 chromosome location (Last three digits of coordinates for cluster start and closer end of interaction from Table 2). **B.** ENSEMBL annotated *PTBP1* transcripts, aligned with the FANTOM5 TSS in A. Note, 948.1 extends in the 5’ direction, to the left beyond the mapped FANTOM5 TSS. **C.** Experimentally determined 5’UTR elements by RLM RACE.

**Figure 3.**
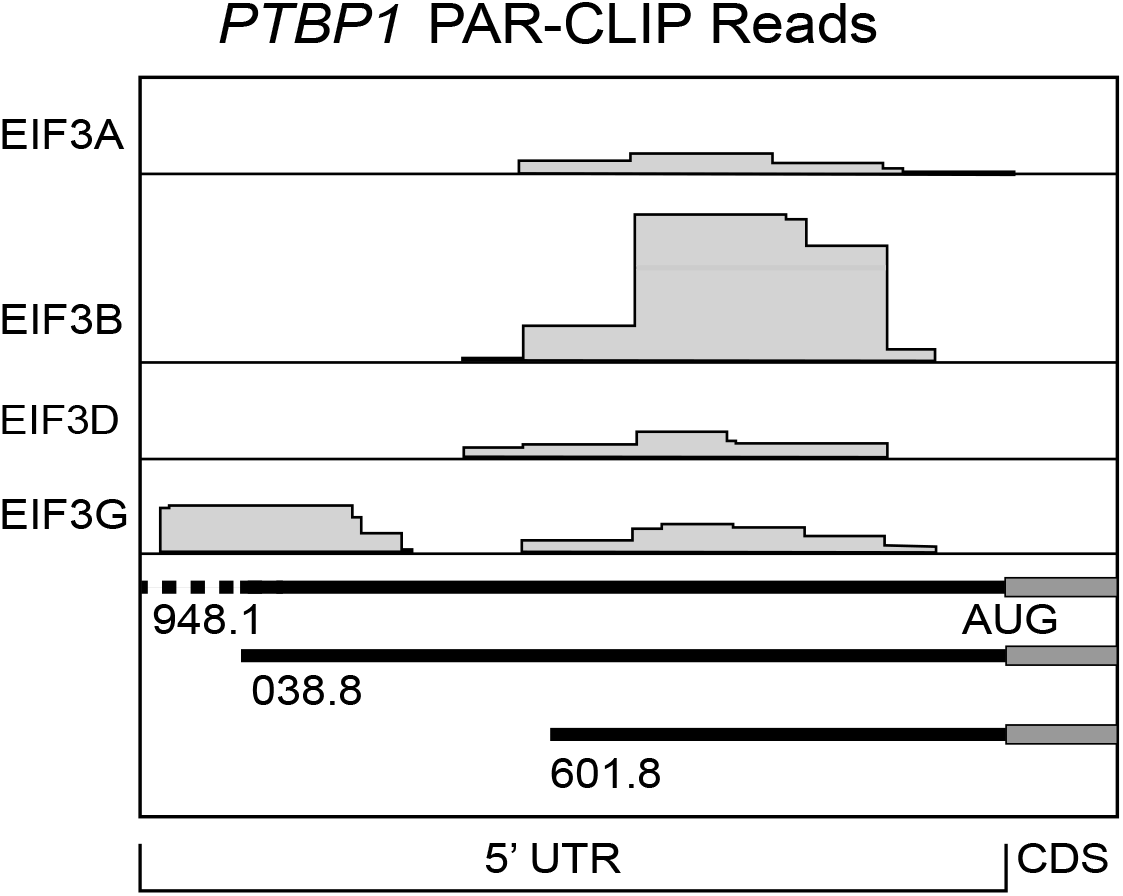
Crosslinking sites between eIF3 and the major *PTBP1* transcript isoforms. Sites of interaction mapped by photoactivatable RNA cross linking and immunoprecipitation (PAR-CLlP) (Lee et al., 2015), by elF3 subunit, indicated to the left. Annotated ENSEMBL transcripts are shown, including the 5’ UTR and the beginning of the CDS, as indicated by the last 4 digits of the ENSEMBL tag (i.e. ENST0000****).

*PTBP1* has three major protein isoforms that only differ with respect to exon 9 inclusion. *PTBP1-1* lacks exon 9 completely, *PTBP1-2* includes only part of exon 9 and *PTBP1-4* contains the full sequence coding for exon 9 (Figure 4A). Although differences in exon properties have been implicated in the different biological roles of PTBP1, the connectivity between the different CDS variants and the mRNA 5’ UTR and 3’ UTR ends is not known. To map the 5’ UTRs for each predicted CDS in the *PTBP1* transcript isoforms, we used a variation of the RLM-RACE methodology (Figure 4B). For each *PTBP1* exon 9 isoform, we observed a single species by RLM-RACE, indicating one major form of 5’ UTR for each CDS variant (Figure 4A and 4C). This was confirmed by a second reaction in which we used a common inner primer to assess the amount of different *PTBP1* 5’ UTRs in the samples (Figure 4D), which revealed two major 5’ UTR species. After sequencing the reactions individually we were able to determine the exact sequence of each transcript up to the cap region. Isoform *PTBP1-1*, which lacks the exon 9 sequence, extends to the 5’ end of the long 5’ UTR, matching the upstream TSS mapped in FANTOM5 (Figure 2A) and the RLM-RACE experiment described above (Figure 2C). By contrast, isoforms *PTBP1-2* and *PTBP1-4* which encode the truncated or full exon 9, respectively, each have the short 5’ UTR, with the downstream TSS mapped in FANTOM5 (Figure 2A and 2C, Figure 4D).

**Figure 4.**
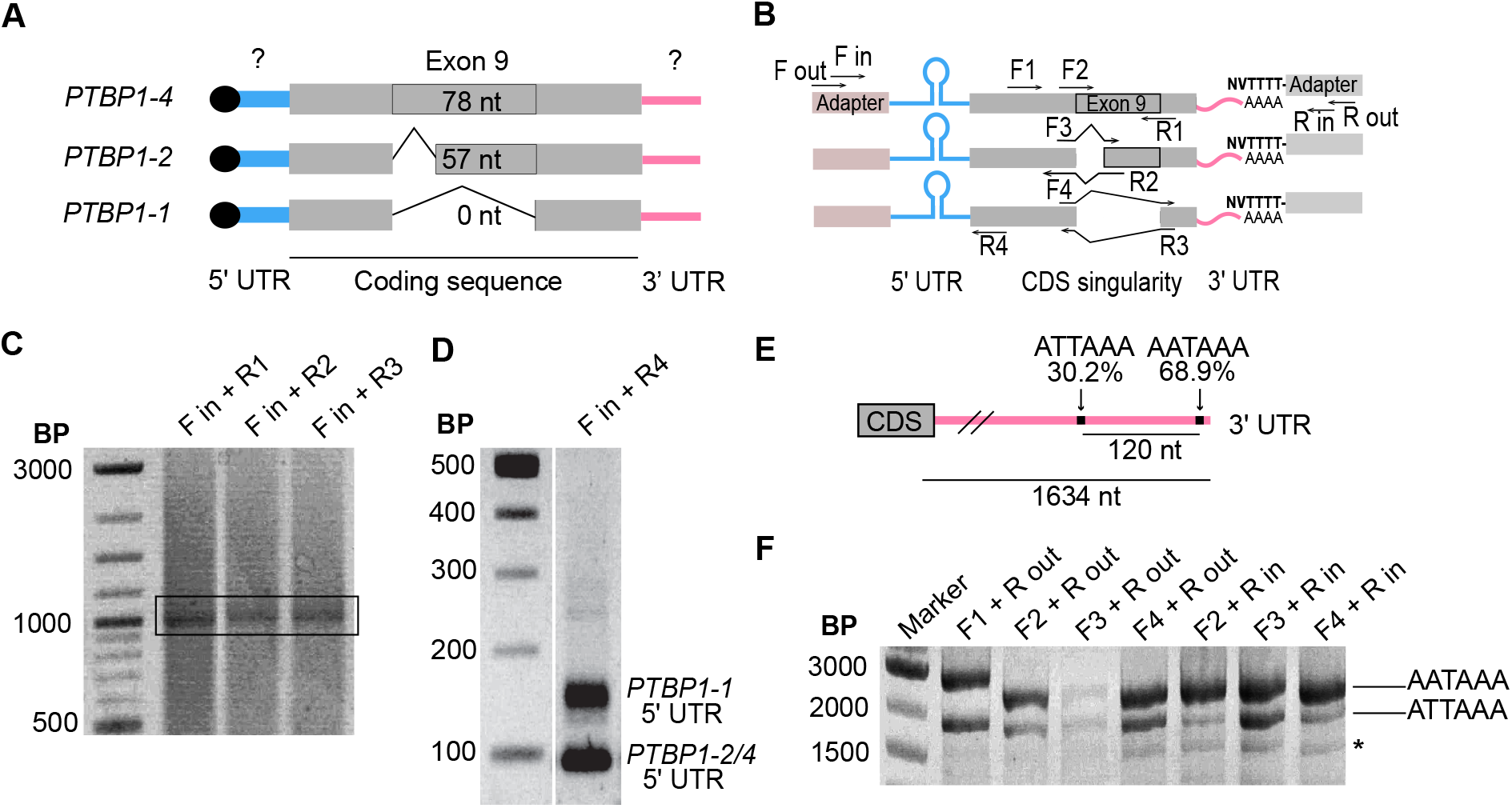
Mapping of the UTR elements of *PTBP1* mRNAs. **A.** Scheme of the composition of the 5’ UTR of *PTBP1* transcripts according to the coding sequence content. **B.** Design of RLM-RACE experiments performed to determine the relationship between 3’ UTR elements, exon 9 boundaries and 5’ UTR elements in *PTBP1* mRNAs. **C.** Agarose gel of final PCR reaction for the 5’ UTR RLM-RACE. Bands in the black rectangle were extracted for sequencing. **D.** 2%agarose gel showing the presence of two known and mapped 5’ UTR lengths during RLM-RACE. **E.** Representation showing high usage polyadenylation sites on *PTBP1* mRNA 3’ UTR (not to scale). Data from (You et al. 2015). **F.** Agarose gel of PCR reactions following RLM-RACE to identify the polyadenylation sites of *PTBP1* transcript isoforms. *Unidentified alternative polyadenylation site.

We also determined the 3’ UTR sequences of *PTBP1* transcript isoforms in HEK239T cells. The APASdb database (You et al. 2015), which contains precise maps and usage quantification of different polyadenylation sites, contains two major polyadenylation sites for *PTBP1* (Figure 4E). We used this information to design specific primers to determine the presence of each polyA site in total RNA extracted from HEK293T cells. By using a forward primer that recognizes the splice junction specific to each transcript upstream of exon 9, we could determine the 3’ UTR length of each isoform by using a reverse primer on a polyA adapter (Figure 4B). The resulting amplification pattern could be visualized by agarose gel (Figure 4F) and then by sequencing. Using this amplification strategy, we observed all three *PTBP1* exon 9 isoforms predicted in the ENSEMBL database to have two different lengths of 3’ UTR resulting from the predicted polyA sites in the APASdb database (Figure 4E and 4F), and possibly a third. Taken together, the present experiments define six *PTBP1* transcript isoforms in HEK293T cells (Figure 5).

**Figure 5.**
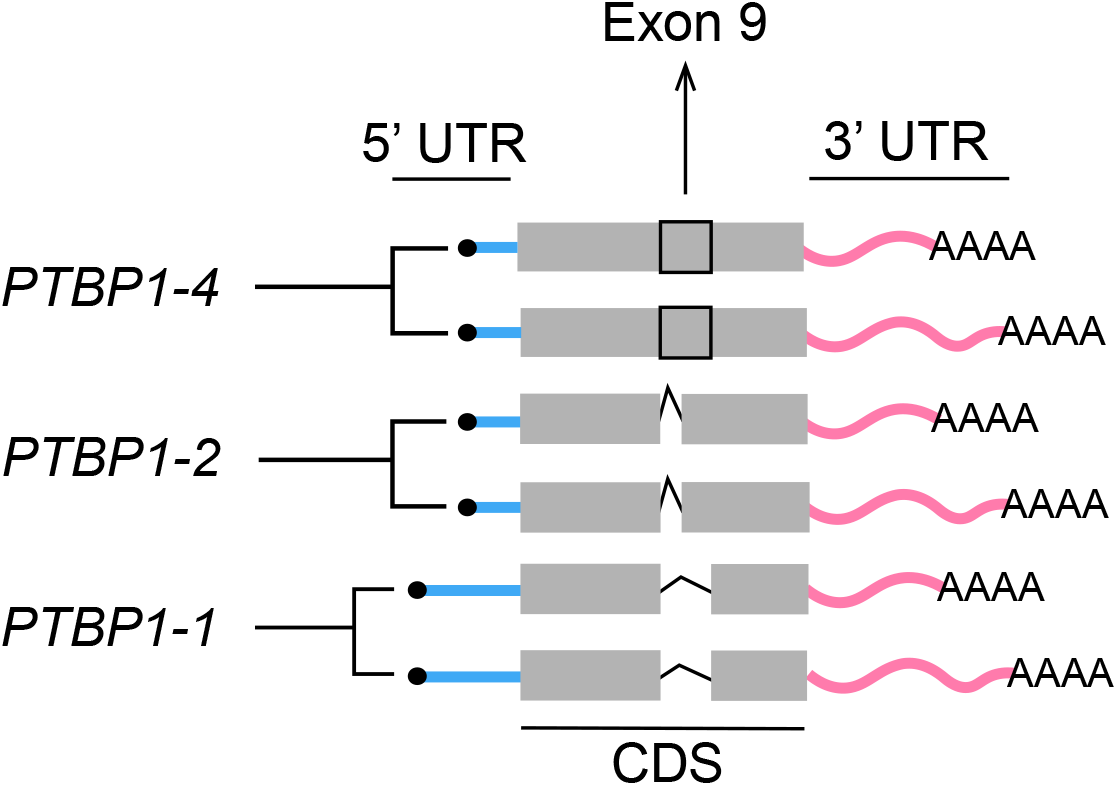
Model for *PTBP1* transcript isoforms in HEK293T cells based on experimental observations. Blue bars, evidence for the existence of two lengths of the 5’ UTR; pink bars, evidence that each transcript has at least two alternative polyadenylation sites, resulting in a long or short 3’ UTR. Grey thick bar represents the alternatively spliced isoforms involving exon 9.

### *PTBP1* 5’ UTR and 3’ UTR contributions to translation regulation

In order to assess whether the differences in PTBP1 expression through the cell cycle are related to the 5’ UTRs and 3’ UTRs in *PTBP1* mRNAs, we used *Renilla* luciferase reporter mRNAs with the different *PTBP1* 5’ UTR and 3’ UTR elements in cell based assays. Using transfections of reporter mRNAs, we first assessed the relative translation levels of each reporter with respect to the cell cycle (Figure 6, Figure 7A). We used 6 hour transfections, as previous results have indicated that these early time points are in the linear range for mRNA transfections (Bert 2006). During the G2 and M phases of the cell cycle, the reporter transcript with the long *PTBP1* 5’ UTR and short *PTBP1* 3’ UTR (Figure 7A) had the highest translation efficiency (Figure 6D and 6E, Figure 7). During the G1 and S phases of the cell cycle, the reporter transcript with the long *PTBP1* 5’ UTR and the long *PTBP1* 3’ UTR had higher translation efficiency (Figure 6B and 6C). Although these experiments are not normalized across cell cycle phases, due to the fact each experiment was carried out separately, we found the experiments synchronized in the G2 and M phases correlated well with unsynchronized cells (Figure 7B and 7C). Furthermore, translation of the reporter mRNAs in the G2 and M phases also correlate well with each other (Figure 7D). By contrast, translation in G1 and S synchronized cells did not correlate with the unsynchronized cells but rather correlated with one another (Figure 8). These results indicate that translation in the G2 and M phases, even though relatively short time-wise (~2 hours total) with respect to the entire cell cycle, dominate translation of the reporter mRNAs with *PTBP1* 5’ UTR and 3’ UTR elements. These results are consistent with endogenous PTBP1 levels observed by Western blotting (Figure 1), suggesting that posttranscriptional regulation of PTBP1 levels occurs to a significant extent at the level of translation during the G2 and M phases of the cell cycle.

**Figure 6.**
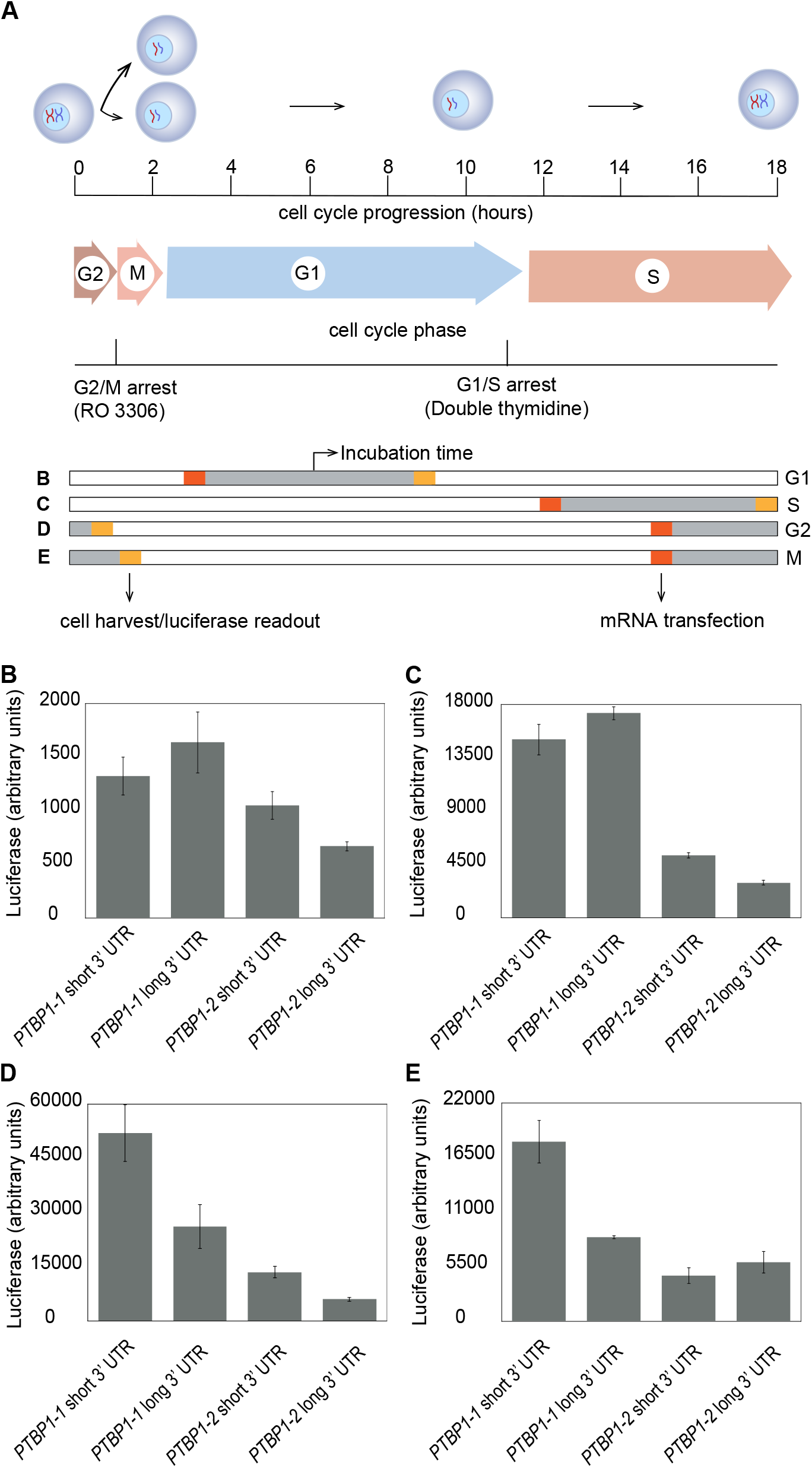
Differences in translation profiles across the cell cycle. **A.** Scheme of mRNA transfection and cell collection on HEK293T cells that were synchronized using RO3306 and double thymidine. Lower schematic, bold letters refer to experiments shown in panels **B** through **E. B.** Luciferase reporter activity in the G1 phase of the cell cycle. **C.** Luciferase reporters activity in S phase. **D.** Luciferase reporter activity in G2. **E.** Luciferase reporter activity in M phase. The lengths of each cell phase were determined experimentally with the use of FACS.

**Figure 7.**
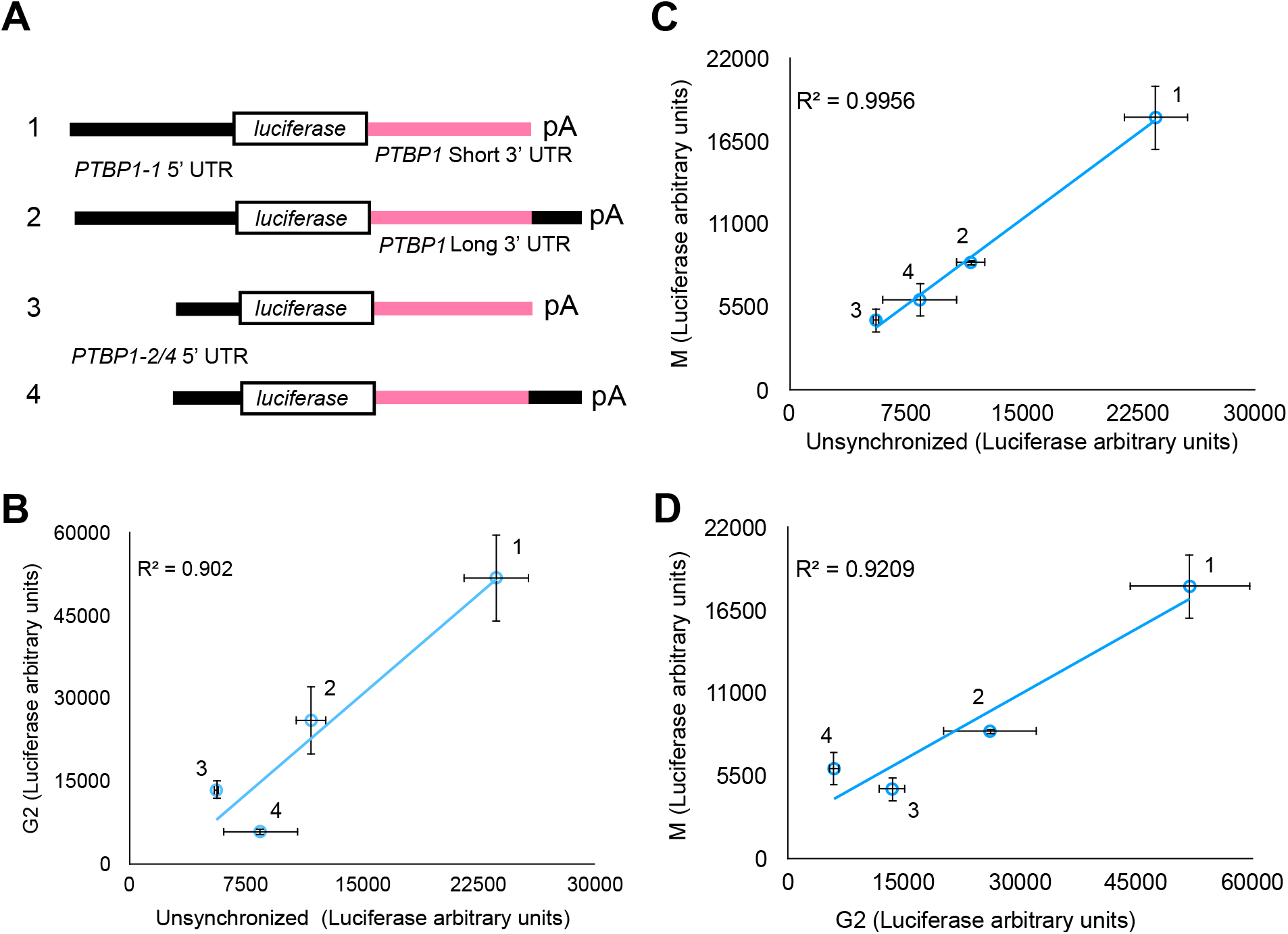
Translation profile of luciferase reporters with *PTBP1* 5’ UTR and 3’ UTR elements in G2 and M phases. **A.** Schematic of luciferase reporters transfected in HEK293T cells. **B.** Luciferase activity from transcripts during G2 plotted against a mixed population of untreated cells. **C.** Luciferase activity from transcripts in M phase plotted against transcripts in a mixed population of untreated cells. **D.** Luciferase activity from transcripts during G2 phase plotted against M phase cells. Transcripts were numbered according to the schematic in A.

**Figure 8.**
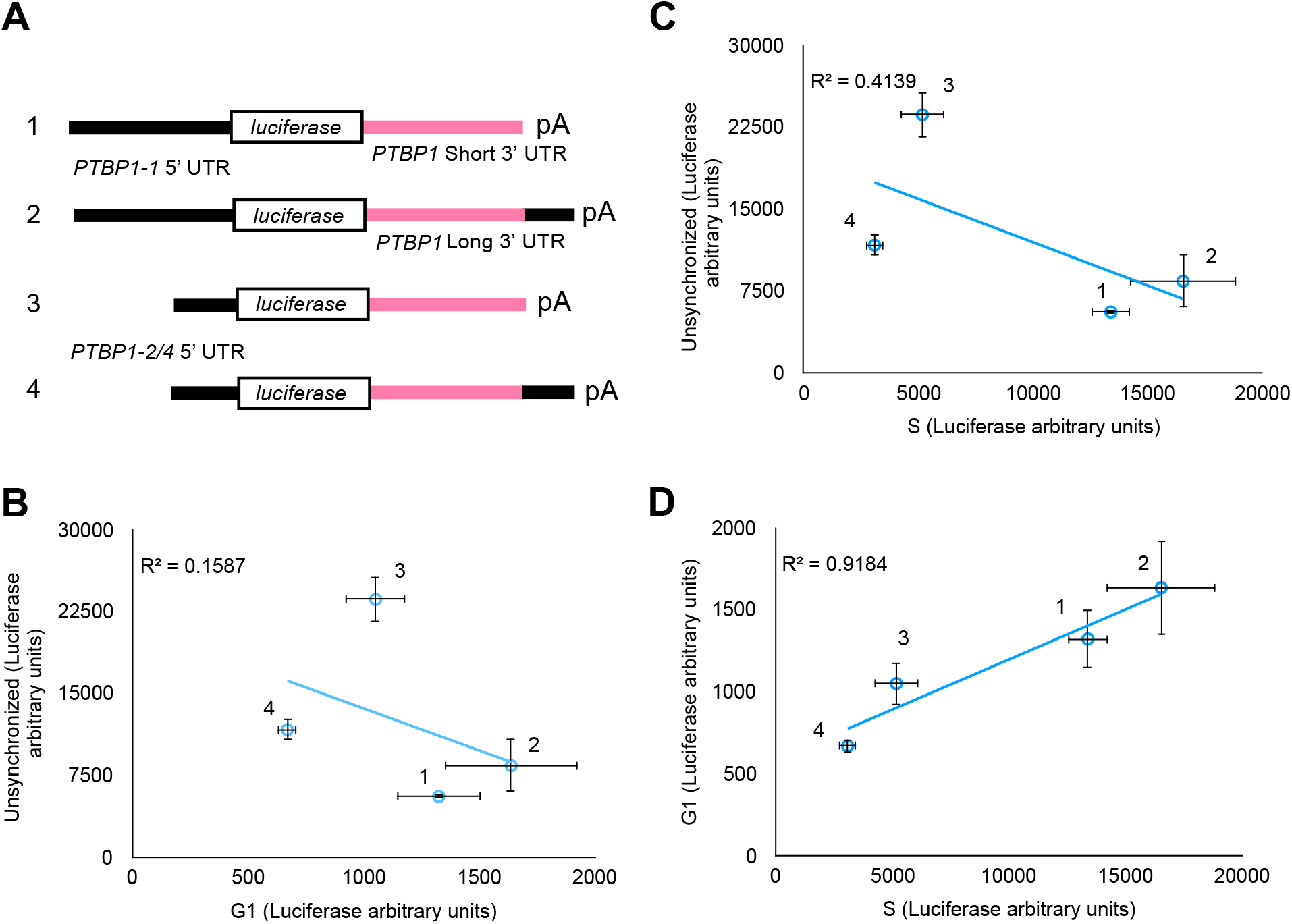
Translation profile of luciferase reporters with *PTBP1* 5’ UTR and 3’ UTR elements in G1 and S phases. **A.** Schematic of the luciferase reporters used for the assay. **B.** Luciferase activity from transcripts during the G1 phase of the cell cycle plotted against a mixed population of cells. **C.** Luciferase activity from transcripts during the S phase plotted against a mixed population of cells. **D.** Luciferase activity from transcripts in G1 plotted against activity from transcripts in the S phase. Transcripts are numbered according to the schematic in A.

### Implications of eIF3 binding to *PTBP1* mRNA on its translation

The results above suggest that translational regulation plays an important role in controlling PTBP1 isoform expression. Given the fact that eIF3 cross-links to specific sequences in the 5’ UTR of *PTBP1* mRNA (Lee et al., 2015), we next tested the importance of these eIF3–5’ UTR interactions in regulating *PTBP1* translation. We used luciferase reporter assays to measure differences in the translation output of mRNAs with the longer *PTBP1* 5’ UTR, which contains two sites of eIF3 crosslinking (Figure 2), or lacking regions known to bind eIF3 (Figure 9A). In untreated HEK293T cells, individually deleting eIF3 crosslinking sites had a minimal impact on translation, whereas deleting both eIF3-interacting regions increased translation of these mRNAs (Figure 9B). In separate experiments, eIF3 immunoprecipitated from cell lysates using an antibody against EIF3B (Lee et al., 2015) was bound to all three endogenous *PTBP1* coding sequence isoforms (Figure 9C).

**Figure 9.**
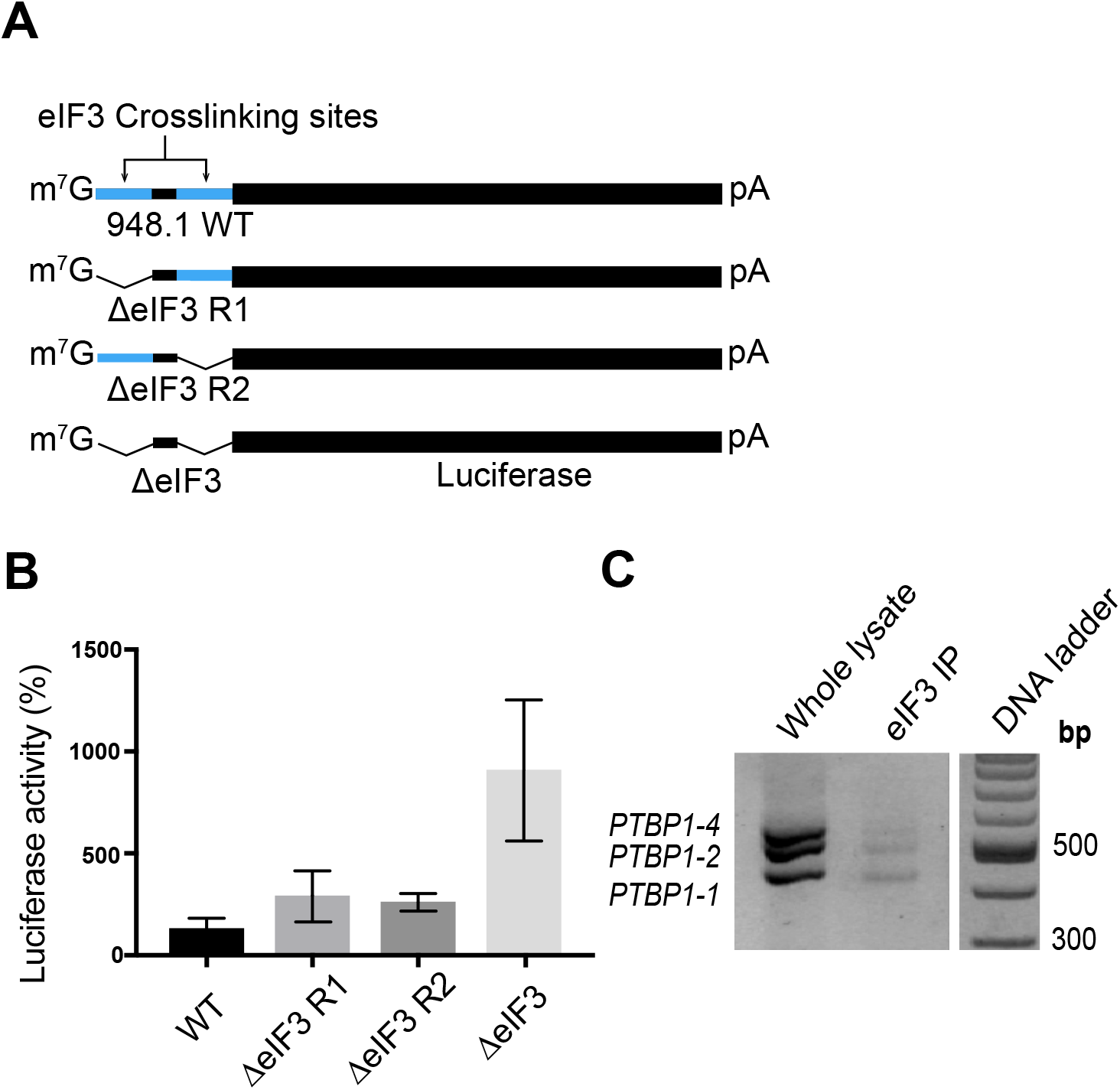
Binding of reporter mRNAs containing sites of interaction with eIF3. **A.** Schematic of *PTBP1* 5’ UTR-luciferase reporter mRNAs. WT, wild type; ΔeIF3, deletion of elF3 PAR-CLIP clusters, nucleotide positions 25-49 (Region 1, R1) and/or 58-86 (Region 2, R2) for the *PTBP1* transcript with the long 5’ UTR (GenBank accession NM_002819). **B.** Luciferase activity in HEK293T cells transfected with mRNAs containing *PTBP1* 5’ UTR elements with or without deletions of the eIF3 cross linking sites in Region 1 (R1) and/or Region 2 (R2). **C.** *PTBP1* mRNA exon 9 coding sequence isoforms that immunoprecipitate with eIF3, as determined by RT-PCR.

To check if the 5’ UTR of *PTBP1* is sufficient for eIF3 binding, and whether both lengths of *PTBP1* 5’ UTR bind similarly to eIF3 across the cell cycle, we designed mRNAs with either the long or short *PTBP1* 5’ UTR sequences upstream of a luciferase open reading frame. We also tested a reporter with a mutated 5’ UTR in which the sequences that cross-link to eIF3 were deleted (Figure 2, Figure 10A). These mRNAs were transfected into HEK293T cells, and the cells were collected in different stages of the cell cycle (Figure 10C). We then immunoprecipitated eIF3 from cell lysates as above (Lee et al., 2015), followed by RNA extraction and quantitative PCR using primers for the luciferase CDS (Figure 10C). Upon deletion of the eIF3 crosslinking sites, eIF3 no longer bound to the reporter mRNAs (Figure 10D). Notably, although the longer *PTBP1* 5’ UTR interacts with eIF3 more efficiently than the short 5’ UTR, both species of 5’ UTR bind to eIF3 more efficiently during the S phase and less so during G2, and even less during G1 (Figure 10D). In the above immunoprecipitation experiments, we used a random 3’ UTR instead of the 3’ UTR elements derived from *PTBP1* transcript isoforms to assess the influence of the *PTBP1* 5’ UTR. To test whether eIF3 binding might also be influenced by the *PTBP1* 3’ UTR, we designed chimeric mRNAs with different combinations of *PTBP1* 5’ UTR and *PTBP1* 3’ UTR, using the same reporter system (Figure 10B). Similarly to the 5’ UTR experiment (Figure 10D), binding of the mRNAs containing the *PTBP1* 3’ UTR to eIF3 is more prevalent during the S phase compared to the other cell phases. Interestingly, the length of the 3’ UTR interacting with eIF3 changes as cell phases progresses, with a switch happening during the mitotic phase (Figure 10E). Altogether, these results indicate that elF3 binds to *PTBP1* mRNAs likely by interacting with both 5’ UTR and 3’ UTR elements in a cell cycle dependent manner (Figure 10).

**Figure 10.**
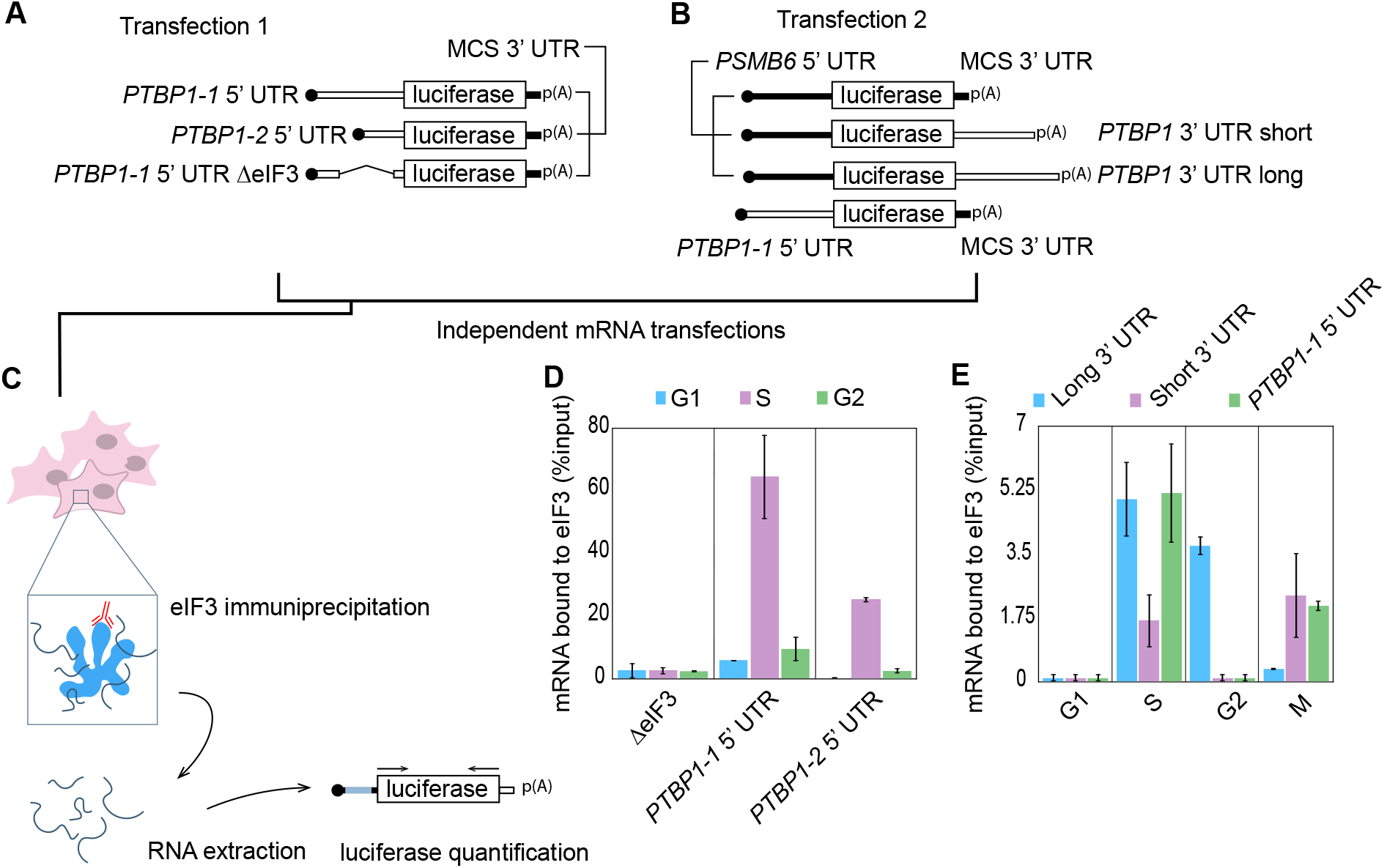
Differential binding of *PTBP1* UTR elements to eIF3 across the cell cycle. **A–B.** Schematics of the luciferase reporters used in the experiments. **C.** Schematic of the transfection, immunoprecipitation and quantification method used to determine luciferase reporter mRNA binding to eIF3. **D.** Distribution of binding to elF3 across the cell cycle for the different *PTBP1* 5’ UTR elements as well as the deletion mutant. **E.** Distribution of binding to eIF3 across the cell cycle for the the different *PTBP1* 3’ UTR elements, as well as the long form of the *PTBP1* 5’ UTR.

## DISCUSSION

A transcript set is the collection of mRNA isoforms that originate from a given genomic sequence. Transcripts are defined by introns, exons, UTRs and their positions. Human transcript set information is stored in large databases and browsers such as ENSEMBL, REFSEQ and UCSC (Zhao and Zhang 2015). However, cases in which the annotations of isoforms are inconsistent across databases are not uncommon (Promponas et al. 2015; Brenner 1999; Schnoes et al. 2009). Given the existence of overlapping, variable transcript isoforms, determining the functional impact of the transcriptome requires identification of full-length transcripts, rather than just the genomic regions that are transcribed (Pelechano et al. 2013). While working with *PTBP1* mRNAs we noticed that sequences available in the ENSEMBL and FANTOM5 databases had discrepancies with respect to the transcription start sites of the major mRNA transcripts (Figure 2). We therefore decided to validate the major mRNA isoforms for *PTBP1* as the basis for future functional analysis of post-transcriptional regulation of PTBP1 expression. We were able to confirm at least six mRNA forms (Figure 5). These mRNA isoforms had differences in the 5’ UTR, coding sequence and 3’ UTR, suggesting that PTBP1 protein isoform expression may be regulated in multiple ways. PTBP1 is a pleiotropic protein, functioning in a variety of cellular processes. It is still unclear if the multiple activities of PTBP1 share a mechanistic pathway and more importantly how PTBP1 could act in posttranscriptional regulation in a tissue-specific way that is singular to the physiology of a certain set of cells. Although PTBP1 has been extensively studied, the multiple *PTBP1* transcript isoforms we have identified will now enable biochemical analysis of *PTBP1* mRNA regulation and function in different stages of the cell cycle.

Identifying RNA exon-exon connectivity remains a challenge when dealing with unknown mRNA isoforms. By combining long read sequencing, RNA-seq and biochemical validation, we were able to fully characterize *PTPB1* transcript isoforms. We used nanopore long read sequencing with the goal to resolve connectivity between 5’ UTR, CDS and 3’ UTR elements of *PTBP1*mRNAs. However, due to the inability of long-read sequencing to accurately reach the 5’ end of mRNA transcripts (Workman et al. 2018), we complemented nanopore sequencing with RNA Ligase Mediated Rapid Amplification of cDNA Ends (RLM-RACE) in order to determine the full length of *PTBP1* mRNA isoforms present in HEK293T cells. This approach should be useful to identify the collection of PTBP1 variants in different cell types or culture conditions (Lundberg et al. 2010). *PTBP1* mRNA has three major isoforms in the coding sequence that differ from each other at exon 9. *PTBP1-1* lacks exon 9, *PTBP1-2* has a partial sequence of exon 9 and *PTBP1-4* has full length exon 9. Because there are three different coding sequences (CDS) (Figure 4A), resulting in three different proteins, and two distinct lengths of 5’ UTR, we aimed at determining the exact full length sequence of each transcript. We found that *PTBP1-1* bears the longer 5’ UTR and *PTBP1-2* and *PTBP1-4* both bear the shorter 5’ UTR. There is only one visible band in the agarose gel for each transcript, meaning that there is only one major form of the 5’ UTR for each transcript (Figure 4C). Consistent with the APASdb database for alternative polyadenylation sites (You et al. 2015), we identified two alternative polyadenylation sites with significant usage, resulting in each of the three major *PTBP1* transcripts having two distinct 3’ UTR lengths (Figure 4E and 4F).

The mapping of all major *PTBP1* transcripts in HEK293T cells (Figure 5) generated the information necessary for the biochemical analysis of *PTBP1* translational regulation. Translational control elements can be located within the 5’ UTR and the 3’ UTR, with overall translation being affected by characteristics such as length, start-site consensus sequences as well as the presence of secondary structure, upstream AUGs, upstream open reading frames (uORFs) and internal ribosome entry sites (IRESs), and binding sites for trans-acting factors (Wilkie et al. 2003). UTR elements have been found to be involved in regulating cell cycle dependent translation. For example, histone translational control in Leishmania requires both 5’ and 3’ UTRs to properly restrict H2A translation to the S phase (Abanades et al. 2009). Differences in 3’ UTR length due to alternative polyadenylation have also been shown to result in acceleration of the cell cycle in cancer cells (Wang et al. 2018). We found that differences in the length of *PTBP1* UTRs results in altered translational efficiency as the cell cycle progresses (Figures 6-8), which may reflect the need to regulate PTBP1 protein isoform translation quickly depending on cellular demands (Sonenberg 1994;Pesole et al. 2001; Mayr 2017).

We previously found that eIF3 binds to PTBP1 through its 5’ UTR (Lee et al.2015) (Figures 2 and 3). Here we found both lengths of 5’ UTR are able to bind to eIF3 through two different sequence regions (Figure 10). We also found that eIF3 can bind to the *PTBP1* 3’ UTRs (Figure 10E). Interestingly, the different UTR lengths have differing impacts on translation (Figures 6-8) and eIF3 binding (Figure 9) in a cell cycle dependent manner. Binding of eIF3 to *PTBP1* mRNA isoforms is most abundant during the S and G2 phases, with the length of the 3’ UTR seeming to influence the extent of eIF3 binding. During S phase, binding is mediated predominantly through the long 3’ UTR and through the 5’ UTR, which correlates with overall repression of translation (Figures 1, 6-10). During G2, eIF3 interacts only with mRNAs bearing the long 3’ UTR, which correlates with repression of translation of these transcripts (Figure 6D, 7B and 7D). Consistent with eIF3 acting as a repressor, previous observations indicate that long 3’ UTRs often repress translation (Szostak and Gebauer 2013; Yamashita and Takeuchi 2017). Future experiments will be required to establish a mechanistic basis for eIF3 repression of *PTBP1* mRNA translation in the S and G2 phases of the cell cycle.

Recently, PTBP1 has been found to be important for cell cycle progression. For example, PTBP1 enables germinal center B cells to progress through the late S phase of the cell cycle rapidly (Monzon-Casanova et al., 2018). In addition, knockout of Ptbp1 in mice results in embryonic lethality due to prolonged G2 to M progression (Shibayama et al. 2009). Notably, we find that PTBP1 expression is highest during the G2 and M cell cycle phases (Figure 1B), which could be explained by the increase in the cell’s demand for PTBP1 in late S phase for proper cell progression. Since the levels of *PTBP1* mRNA during the different stages of the cell cycle remained relatively unchanged (Figure 1C), post-transcriptional regulation seems to be central to PTBP1 expression. With curated information on *PTBP1* mRNA isoforms present in HEK293 mammalian cells (Figure 5), it will now be possible to dissect the post-transcriptional regulatory mechanisms involved in cell cycle dependent expression of PTBP1 isoforms, and the downstream physiological consequences.

## MATERIALS AND METHODS

### Cells and transfections

Human HEK293T cells were cultured in DMEM (Invitrogen) supplemented with 10% of Fetal Bovine Serum (FBS) (Seradigm) and 1% Pen/Strep (Gibco, Cat. # 15140122). RNA transfections were performed using MirusTranslT^®^-mRNA Transfection Kit (Cat. # MIR 2250), with the following modifications to the manufacturer’s protocol. Sixteen hours before transfection, HEK293T cells were seeded into opaque 96-well plates to reach ~80% confluence at the time of transfection. For each well, 9 μL of pre-warmed OptiMEM (Invitrogen) was mixed with 250 ng of RNA, 0.27 μL of Boost reagent and 0.27 μL of TransIT mRNA reagent. Reactions were incubated for 3 minutes at room temperature, added drop-wise to the well, and luciferase activity was measured 6–8 hours (as indicated) after transfection, using. using the Renilla Luciferase assay kit (Promega, Cat. # E2820) and a Microplate 309 Luminometer (Veritas).

For G1 transfections and luminescence readouts, cells were grown to 30% confluence and compound RO3306 (Tanenbaum et al. 2015; Vassilev et al. 2006) was added to a 6 μM final concentration. Cells were incubated for 18 hours. After 18 hours, cells were released and incubated for two hours with fresh media. After this time, cells were transfected with the desired mRNA. Luminescence was measured after six hours of incubation. For S transfections and readouts, cells were grown to 20% confluence in standard media and thymidine was added to a final concentration of 2 mM. Cells were incubated for 18 hours in a tissue culture incubator and after 18 hours, thymidine was removed by two washes of HBSS media (Invitrogen). Fresh media was added and cells were incubated for 9 hours. After 9 hours, thymidine was added again to a 2 mM final concentration. Cells were incubated in tissue culture incubator for 15 hours. After 15 hours, cells were washed with HBSS media (Invitrogen) and released into fresh media. After 1 hour of incubation, cells were transfected with desired mRNA and luminescence was measured after 6 hours. For G2 transfection and luminescence readouts, cells were synchronized with the same protocol as for S phase. After release, however, they were incubated for 4 hours before transfection. After transfection, cells were incubated for three hours and RO3306 was added following the previous protocol. This guaranteed the cells would not progress into M phase before luminescence was measured. Cells were then incubated for three hours and luminescence was assessed. For M transfection and readouts, cells were treated exactly as G2, expect RO3306 was not added, allowing them to progress into M phase after four hours. Luminescence was measured after 6 hours of incubation.

### Cell cycle analysis

The cell cycle analysis was performed using propidium iodide (PI) staining to determine the percent of cells in each phase of the cell cycle. HEK293T cells were fixed in 70% ethanol at the time of harvesting and stained with PI. They were treated with ribonuclease, to ensure only DNA would be stained, and 200 μL of PI (50 μg/mL) was added. They were immediately analyzed after staining on a BD Fortessa Flow Cytometer.

### Plasmids

To generate the luciferase plasmids used on this work, sections of either the *PTBP1* 5’ UTR (GenBank accession NM_002819) or the *PTBP1* 3’ UTR were first amplified from human cDNA extracted from HEK293T cells. These were then placed downstream of a T7 RNA polymerase promoter using overlap extension PCR and InFusion cloning. The 5’ UTRs were then inserted together with *Renilla* luciferase into plasmid pcDNA4 V102020 (lnvitrogen). The eIF3 binding mutants and *PSMB6-PTBP1* chimeras were made by insertional mutagenesis with primers annealed to the pcDNA4 plasmid digested at the desired insertion site. Primers and sequences are included in Table 1.

**Table 1.**
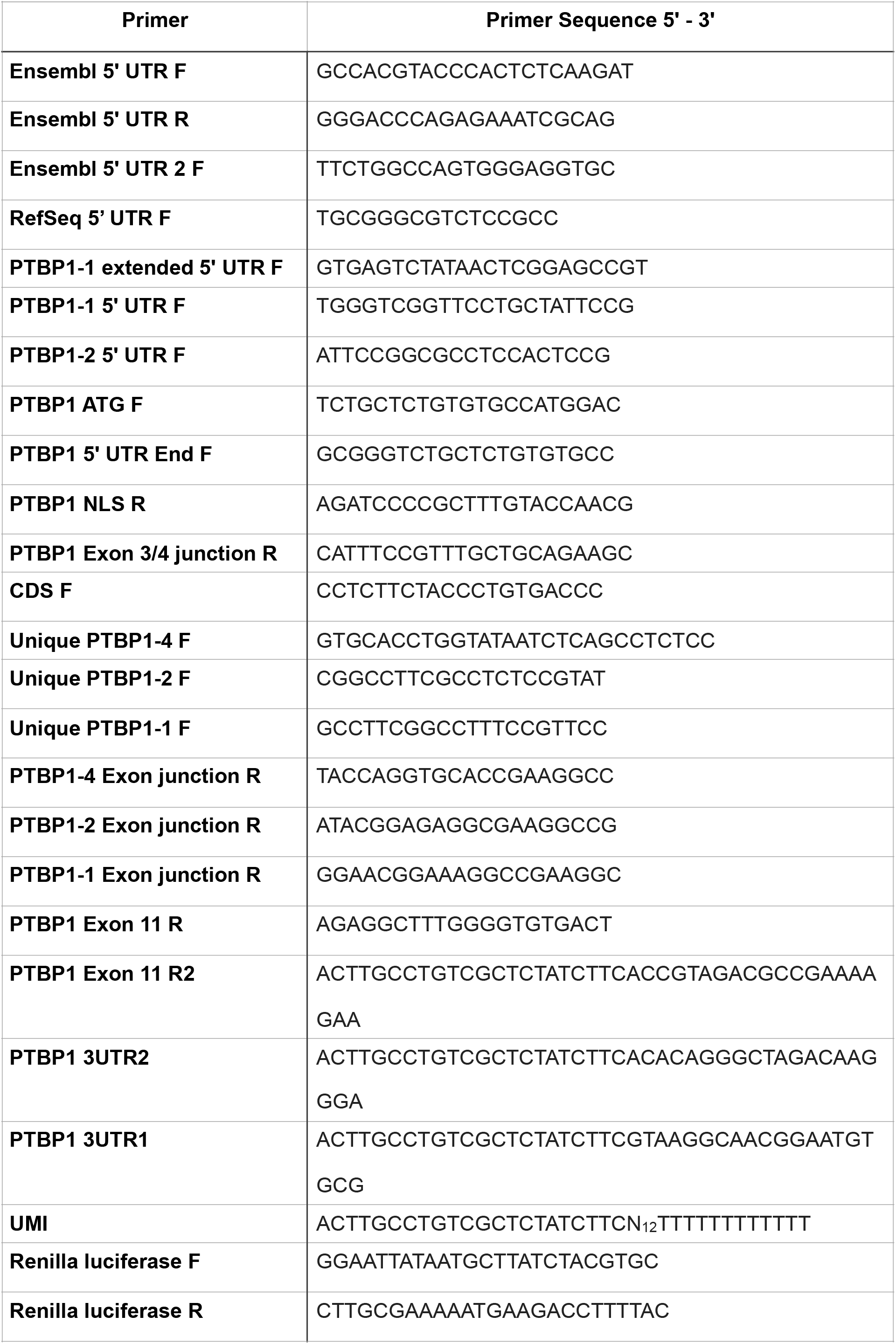
Primers used in this study.

**Table 2.**
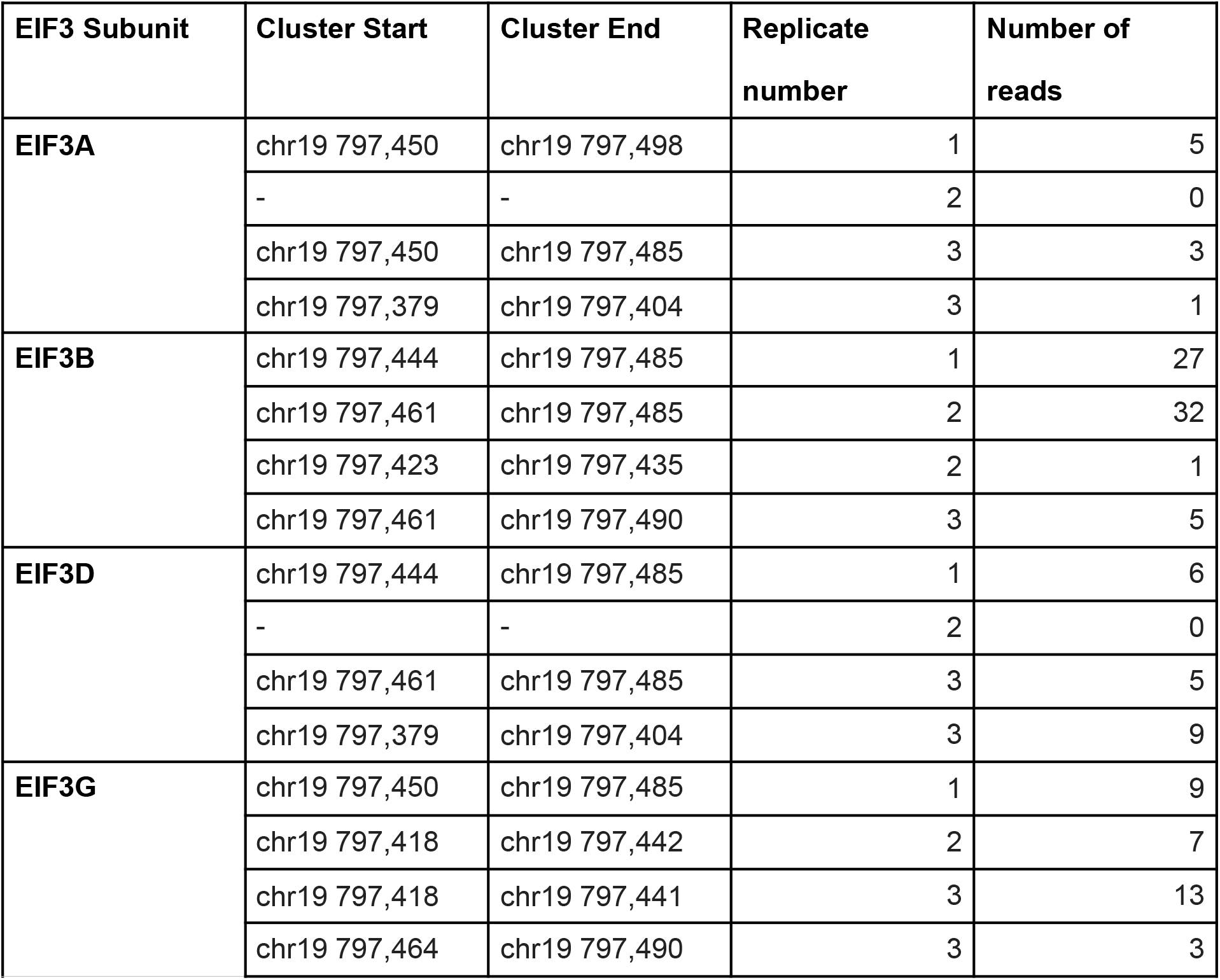
PAR-CLIP crosslinking sites in hg38 coordinates.

| **EIF3 Subunit** | **Cluster Start** | **Cluster End** | **Replicate number** | **Number of reads** |
| --- | --- | --- | --- | --- |
| **EIF3A** | chr19 797,450 | chr19 797,498 | 1 | 5 |
|  | - | - | 2 | 0 |
|  | chr19 797,450 | chr19 797,485 | 3 | 3 |
|  | chr19 797,379 | chr19 797,404 | 3 | 1 |
| **EIF3B** | chr19 797,444 | chr19 797,485 | 1 | 27 |
|  | chr19 797,461 | chr19 797,485 | 2 | 32 |
|  | chr19 797,423 | chr19 797,435 | 2 | 1 |
|  | chr19 797,461 | chr19 797,490 | 3 | 5 |
| **EIF3D** | chr19 797,444 | chr19 797,485 | 1 | 6 |
|  | - | - | 2 | 0 |
|  | chr19 797,461 | chr19 797,485 | 3 | 5 |
|  | chr19 797,379 | chr19 797,404 | 3 | 9 |
| **EIF3G** | chr19 797,450 | chr19 797,485 | 1 | 9 |
|  | chr19 797,418 | chr19 797,442 | 2 | 7 |
|  | chr19 797,418 | chr19 797,441 | 3 | 13 |
|  | chr19 797,464 | chr19 797,490 | 3 | 3 |

### Western Blot

Western Blot analysis was carried out using the following antibodies: anti-ElF3B (Bethyl A301-761A), anti-HSP90 (BD 610418) and anti-PTBP1 (MABE986, clone BB7); all antibodies were used with a 1:10000 dilution.

### *In vitro* transcription

RNAs to be used for transfections were made by *in vitro* transcription with T7 RNA polymerase (NEB). For luciferase mRNAs, transcription was performed in the presence of 3’-O-Me-m^7^G (5’)ppp(5’)G RNA Cap Structure Analogue (NEB), using linearized plasmid as template, then polyadenylated using poly-A polymerase (Invitrogen). RNAs were purified by phenol-chloroform extraction and ethanol precipitation or using the RNA clean and concentrator kit (Zymo Research). RNA quality was verified using 2%agarose gels, to ensure mRNAs were intact before transfection.

### RNA Immunoprecipitation and RT-PCR

HEK293T cells grown on 10 cm plates were lysed as needed in three volumes of NP40 lysis buffer (50 mM HEPES-KOH pH=7.5, 500 mM KCl, 2 mM EDTA, 1%Nonidet P-40 alternative, 0.5 mM DTT). Dynabeads were prepared with rabbit IgG (Cell Signaling 2729) and rabbit anti-EIF3B antibody (Bethyl A301-761A). The lysate was split into three parts, the Dynabeads-antibody mixture was added, and the suspensions incubated for 2 h at 4 ° C. The beads were washed four times with NP40 buffer, and bound RNAs were isolated by phenol chloroform extraction and ethanol precipitation. The resulting cDNA was reverse transcribed using random hexamers and Superscript III (Thermo Fisher scientific), and PCR was performed using DNA polymerase Q5 (NEB).

### Oxford Nanopore sequencing

Nanopore sequencing was carried out using the manufacturer protocol for 1D Strand switching cDNA by ligation (SQK-LSK108). The user defined primer used was specific for exon 11 in *PTBP1* mRNA:

5’-ACTTGCCTGTCGCTCTATCTTCAGAGGCTTTGGGGTGTGACT-3’

### Rapid Amplification of cDNA ends (RACE)

RACE analysis followed the protocol described for the FirstChoice RLM-RACE kit (Ambion), using the thermostable Vent DNA polymerase (NEB) and the adapter primers provided by the kit. The user defined primers were:

PTBP1-2 Exon junction reverse (R2): 5’-ATA CGG AGA GGC GAA GGC CG-3’
PTBP1-1 Exon junction reverse (R3): 5’-GGA ACG GAA AGG CCG AAG GC-3’
PTBP1-4 Exon junction reverse (R1): 5’-TAC CAG GTG CAC CGA AGG CC-3’
PTBP1 general reverse (R4): 5’-AGA TCC CCG CTT TGT ACC AAC G-3’

For the 3’ UTR RACE the user defined primers were:

Unique PTBP1-4 (F2): 5’-GTGCACCTGGTATAATCTCAGCCTCTCC-3’
Unique PTBP1-2 (F3): 5’-CGGCCTTCGCCTCTCCGTAT-3’
Unique PTBP1-1 (F4): 5’-GCCTTCGGCCTTTCCGTTCC-3’
CDS F (F1): 5’-CCTCTTCTACCCTGTGACCC-3’

## ACKNOWLEDGMENTS

We thank D. Black for providing the PTBP1 antibody (Bb7) and for helpful discussions. We also thank W. Li, F. R. Ward, and A. S.-Y. Lee for helpful discussions. This work was funded by a predoctoral fellowship to L.M.A.T. through CAPES Science Without Borders (fellowship P-3-03822) and by grant P50-GM102706 from NIGMS to J.H.D.C.

## REFERENCES

Abanades DR, Ramírez L, Iborra S, Soteriadou K, González VM, Bonay P, Alonso C, Soto M. 2009. Key role of the 3’ untranslated region in the cell cycle regulated expression of the Leishmania infantum histone H2A genes: minor synergistic effect of the 5’ untranslated region. BMC Mol Biol 10: 48.

Bert AG. 2006. Assessing IRES activity in the HIF-1 and other cellular 5’ UTRs. RNA 12: 1074–1083.

Brenner SE. 1999. Errors in genome annotation. Trends Genet 15: 132–133.

Garcia-Blanco MA, Jamison SF, Sharp PA. 1989. Identification and purification of a 62,000-dalton protein that binds specifically to the polypyrimidine tract of introns. Genes Dev 3: 1874–1886.

Gil A, Sharp PA, Jamison SF, Garcia-Blanco MA. 1991. Characterization of cDNAs encoding the polypyrimidine tract-binding protein. Genes Dev 5: 1224–1236.

Gueroussov S, Gonatopoulos-Pournatzis T, Irimia M, Raj B, Lin Z-Y, Gingras A-C, Blencowe BJ. 2015. An alternative splicing event amplifies evolutionary differences between vertebrates. Science 349: 868–873.

Hinnebusch AG, Ivanov IP, Sonenberg N. 2016. Translational control by 5’-untranslated regions of eukaryotic mRNAs. Science 352: 1413–1416.

Kamath RV, Leary DJ, Huang S. 2001. Nucleocytoplasmic shuttling of polypyrimidine tract-binding protein is uncoupled from RNA export. Mol Biol Cell 12: 3808–3820.

Lee ASY, Kranzusch PJ, Cate JHD. 2015. eIF3 targets cell-proliferation messenger RNAs for translational activation or repression. Nature 522: 111–114.

Li B, Yen TSB. 2002. Characterization of the nuclear export signal of polypyrimidine tract-binding protein. J Biol Chem 277: 10306–10314.

Lundberg E, Fagerberg L, Klevebring D, Matic I, Geiger T, Cox J, Algenäs C, Lundeberg J, Mann M, Uhlen M. 2010. Defining the transcriptome and proteome in three functionally different human cell lines. Mol Syst Biol 6: 450.

Ma W, Mayr C. 2018. A Membraneless Organelle Associated with the Endoplasmic Reticulum Enables 3′UTR-Mediated Protein-Protein Interactions. Cell. http://dx.doi.org/10.1016/j.cell.2018.10.007.

Mayr C. 2017. Regulation by 3’-Untranslated Regions. Annu Rev Genet 51: 171–194.

Monzón-Casanova E, Screen M, Díaz-Muñoz MD, Coulson RMR, Bell SE, Lamers G, Solimena M, Smith CWJ, Turner M. 2018. The RNA-binding protein PTBP1 is necessary for B cell selection in germinal centers. Nat Immunol 19: 267–278.

Noguchi S, Arakawa T, Fukuda S, Furuno M, Hasegawa A, Hori F, Ishikawa-Kato S, Kaida K, Kaiho A, Kanamori-Katayama M, et al. 2017. FANTOM5 CAGE profiles of human and mouse samples. Sci Data 4: 170112.

O’Leary NA, Wright MW, Rodney Brister J, Ciufo S, Haddad D, McVeigh R, Rajput B, Robbertse B, Smith-White B, Ako-Adjei D, et al. 2015. Reference sequence (RefSeq) database at NCBI: current status, taxonomic expansion, and functional annotation. Nucleic Acids Res 44: D733–D745.

Pelechano V, Wei W, Steinmetz LM. 2013. Extensive transcriptional heterogeneity revealed by isoform profiling. Nature 497: 127–131.

Pérez I, McAfee JG, Patton JG. 1997. Multiple RRMs Contribute to RNA Binding Specificity and Affinity for Polypyrimidine Tract Binding Protein†. Biochemistry 36: 11881–11890.

Pesole G, Mignone F, Gissi C, Grillo G, Licciulli F, Liuni S. 2001. Structural and functional features of eukaryotic mRNA untranslated regions. Gene 276: 73–81.

Promponas VJ, Iliopoulos I, Ouzounis CA. 2015. Annotation inconsistencies beyond sequence similarity-based function prediction - phylogeny and genome structure. Stand Genomic Sci 10: 108.

Pyo C-W, Choi JH, Oh S-M, Choi S-Y. 2013. Oxidative stress-induced cyclin D1 depletion and its role in cell cycle processing. Biochim Biophys Acta 1830: 5316–5325.

Romanelli MG, Diani E, Lievens PM-J. 2013. New insights into functional roles of the polypyrimidine tract-binding protein. Int J Mol Sci 14: 22906–22932.

Sawicka K, Bushell M, Spriggs KA, Willis AE. 2008. Polypyrimidine-tract-binding protein: a multifunctional RNA-binding protein. Biochem Soc Trans 36: 641–647.

Schnoes AM, Brown SD, Dodevski I, Babbitt PC. 2009. Annotation error in public databases: misannotation of molecular function in enzyme superfamilies. PLoS Comput Biol 5: e1000605.

Shibayama M, Ohno S, Osaka T, Sakamoto R, Tokunaga A, Nakatake Y, Sato M, Yoshida N. 2009. Polypyrimidine tract-binding protein is essential for early mouse development and embryonic stem cell proliferation. FEBS J 276: 6658–6668.

Sonenberg N. 1994. mRNA translation: influence of the 5′ and 3′ untranslated regions. Curr Opin Genet Dev 4: 310–315.

Szostak E, Gebauer F. 2013. Translational control by 3’-UTR-binding proteins. Brief Funct Genomics 12: 58–65.

Tanenbaum ME, Stern-Ginossar N, Weissman JS, Vale RD. 2015. Regulation of mRNA translation during mitosis. Elife 4. http://dx.doi.org/10.7554/elife.07957.

Valcárcel J, Gebauer F. 1997. Post-transcriptional regulation: the dawn of PTB. Curr Biol 7: R705–8.

Vassilev LT, Tovar C, Chen S, Knezevic D, Zhao X, Sun H, Heimbrook DC, Chen L. 2006. Selective small-molecule inhibitor reveals critical mitotic functions of human CDK1. Proc Natl Acad Sci U S A 103: 10660–10665.

Wang Q, He G, Hou M, Chen L, Chen S, Xu A, Fu Y. 2018. Cell Cycle Regulation by Alternative Polyadenylation of CCND1. Sci Rep 8: 6824.

Wang X, He X, Deng X, He Y, Zhou X. 2017. Roles of miR-4463 in H2O2-induced oxidative stress in human umbilical vein endothelial cells. Mol Med Rep 16: 3242–3252.

Wilkie GS, Dickson KS, Gray NK. 2003. Regulation of mRNA translation by 5’- and 3’- UTR-binding factors. Trends Biochem Sci 28: 182–188.

Wollerton MC, Gooding C, Robinson F, Brown EC, Jackson RJ, Smith CW. 2001. Differential alternative splicing activity of isoforms of polypyrimidine tract binding protein (PTB). RNA 7: 819–832.

Wollerton MC, Gooding C, Wagner EJ, Garcia-Blanco MA, Smith CWJ. 2004. Autoregulation of polypyrimidine tract binding protein by alternative splicing leading to nonsense-mediated decay. Mol Cell 13: 91–100.

Workman RE, Tang A, Tang PS, Jain M, Tyson JR, Zuzarte PC, Gilpatrick T, Razaghi R, Quick J, Sadowski N, et al. 2018. Nanopore native RNA sequencing of a human poly(A) transcriptome. http://dx.doi.org/10.1101/459529.

Xue Y, Zhou Y, Wu T, Zhu T, Ji X, Kwon Y-S, Zhang C, Yeo G, Black DL, Sun H, et al. 2009. Genome-wide analysis of PTB-RNA interactions reveals a strategy used by the general splicing repressor to modulate exon inclusion or skipping. Mol Cell 36: 996–1006.

Yamashita A, Takeuchi O. 2017. Translational control of mRNAs by 3’-Untranslated region binding proteins. BMB Rep 50: 194–200.

You L, Wu J, Feng Y, Fu Y, Guo Y, Long L, Zhang H, Luan Y, Tian P, Chen L, et al. 2015. APASdb: a database describing alternative poly(A) sites and selection of heterogeneous cleavage sites downstream of poly(A) signals. Nucleic Acids Res 43: D59–67.

Zhao S, Zhang B. 2015. A comprehensive evaluation of ensembl, RefSeq, and UCSC annotations in the context of RNA-seq read mapping and gene quantification. BMC Genomics 16: 97.

